# Widespread purine bias in bacterial genes driven by runaway transcription

**DOI:** 10.1101/2025.08.25.672197

**Authors:** K. Julia Dierksheide, James C. Taggart, Grace E. Johnson, Gene-Wei Li

## Abstract

Genes in many bacteria are rich in purine nucleotides and poor in pyrimidines. We show that this purine preference is critical for gene expression because it prevents premature transcription termination in species that exhibit runaway transcription. In contrast to coupled transcription-translation^1–5^, runaway RNA polymerases that outpace trailing ribosomes have exposed nascent RNA and are vulnerable to the termination factor Rho^6,7^. Using a massively parallel reporter assay in *Bacillus subtilis*, we found that Rho-dependent termination requires a high C-to-G skew and high T content. Consequently, purine-rich coding (sense) sequences escape premature termination, whereas the correspondingly pyrimidine-rich antisense sequences are targeted by Rho and transcriptionally silenced. This purine requirement drives biased codon usage in most bacterial species with runaway transcription, except in lineages that have lost Rho. Our results suggest that the avoidance of premature transcription termination imposes major constraints on nucleotide content during genome evolution and adaptation of foreign genes.

---

Protein production requires complete synthesis of mRNA molecules. In the model bacterium *Escherichia coli,* premature termination of mRNA transcription is inhibited by a closely trailing ribosome, which prevents nascent RNAs from triggering either intrinsic or Rho-dependent termination^1,4,5,8,9^. However, in other bacteria, transcription is not always tightly coupled to translation: RNA polymerases (RNAPs) outpace ribosomes in many low-GC-gram-positives, such as *Bacillus subtilis*^6,7^. It remains unclear what alternative mechanisms broadly protect these “runaway” RNAPs from premature termination.

In particular, runaway RNAPs are vulnerable to the conserved transcription termination factor Rho. In bacteria, Rho is thought to function as a global surveillance factor by terminating untranslated RNAs^9–13^—inhibition of Rho in both *E. coli* and *B. subtilis* dramatically elevates antisense, but not sense, transcription^14–17^. In *E. coli*, Rho recognizes a series of up to six CC or UC dinucleotides interspersed on nascent RNA^18–21^. This signal is degenerate enough that many RNAs can be targeted in the absence of a closely trailing ribosome^1,5,8,11–13,22–25^. Indeed, in the classic examples of polarity, premature stop codons frequently result in Rho-dependent termination of downstream transcription^9,26–30^. However, for species where runaway RNAPs expose nascent mRNAs to Rho, this promiscuous targeting model would predict widespread termination and is not compatible with the observation of selective termination of antisense transcription.

Based on an observation that *B. subtilis* genes are particularly rich in purines (A and G), complementing the degenerate pyrimidine preference (C and U) of *E. coli* Rho, we hypothesized that sequence, rather than translation state, may form the basis for Rho’s selective avoidance of mRNA in *B. subtilis.* Using a large-scale reporter library to identify regions of transcription termination across the *B. subtilis* genome, we show that sequence composition alone allows *B. subtilis* genes to escape Rho termination. In contrast to *E. coli,* we found that *B. subtilis* Rho activity is highly sequence-specific, only terminating transcription of a small set of genomic fragments strongly biased towards antisense regions with high C and U content. Avoidance of such pyrimidine-rich regions appears to broadly drive purine bias in genes and constrain codon usage across Bacilli that encode Rho. This global coding constraint not only shapes gene evolution within individual species but also influences foreign gene expression. More broadly, our study highlights the outsized influence of a single transcription factor on the sequence of genes, cracking a hidden code within bacterial genomes.

## Genome-wide mapping of Rho termination

If sequence composition protects mRNA from Rho termination in *B. subtilis,* then isolated, untranslated fragments of coding sequence should not be terminated by Rho. To test this hypothesis, we designed a massively parallel reporter assay to measure Rho termination during transcription of 10^5^ DNA fragments (mean length = 286 bp, SD = 74 bp) tiling the *B. subtilis* genome (5.4-fold coverage over 98% of the genome). We linked transcription termination to chloramphenicol resistance via a transcriptional reporter library where DNA fragments are placed upstream of *lacI*, which represses expression of the resistance gene (Fig. 1a). Termination within a fragment reduces *lacI* expression, activating resistance and allowing growth in chloramphenicol. After chloramphenicol selection of the pooled reporter library, variant enrichment is quantified using deep DNA sequencing, revealing fragments that carry termination signals. Rho-dependent termination events were identified by a loss of enrichment when the selection was carried out in a separate culture containing the Rho inhibitor bicyclomycin-benzoate^31^.

**Fig. 1:**
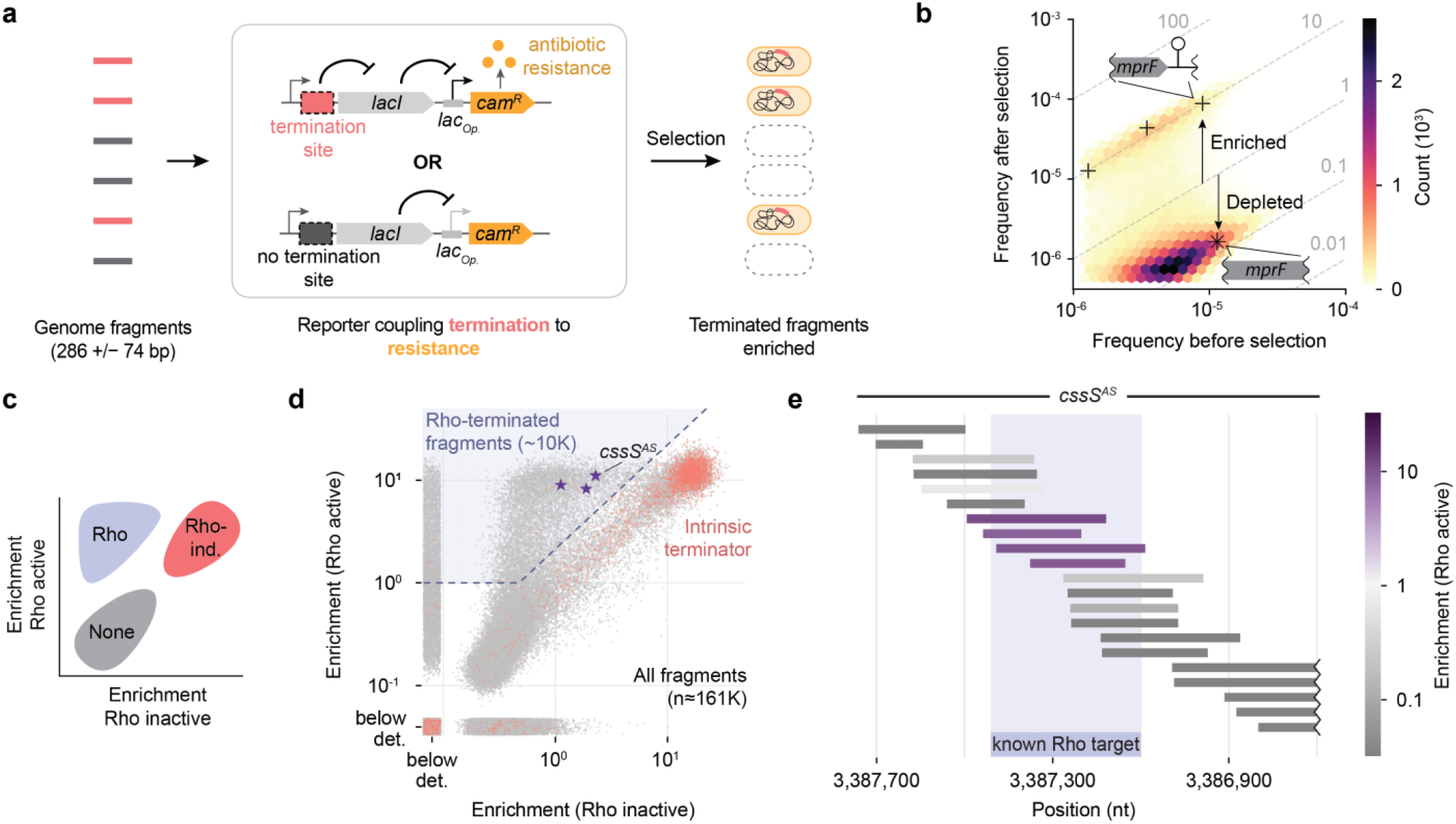
Genome-wide screen for sequences subject to Rho-dependent termination. **a**, Schematic of growth-based massively parallel reporter assay for termination activity. Left, genomic DNA fragments containing (pink) and lacking (grey) transcription termination signals. Mean fragment length was 286 bp with a standard deviation of 74 bp. Middle, reporter coupling termination to antibiotic resistance. *lacI*, lac repressor gene; *lac_Op_*, lac operator; *cam^R^*, chloramphenicol resistance gene; orange circles, protein conferring antibiotic resistance. Selection refers to growth in chloramphenicol. Right, dead (white) and resistant (orange) bacterial cells. **b**, Distribution of fragment frequencies before vs. after selection (Rho active; bicyclomycin-benzoate absent). Grey dashed lines mark enrichment values. Daggers indicate fragments that contain the intrinsic terminator following *mprF*^32^. Asterisk indicates fragment completely overlapping with the *mrpF* coding strand. **c**, Schematic indicating expected positions of fragments with no termination (None), Rho-dependent termination (Rho), or Rho-independent termination (Rho-ind.) activity in plot shown in **d**. **d**, Enrichment of each fragment in the presence (Rho inactive) vs. absence (Rho active) of the Rho inhibitor bicyclomycin-benzoate (BCM). Rho-terminated fragments are defined as those in the region above the dashed lines, shaded in purple (Methods). Purple stars indicate putative Rho-terminated fragments antisense to *cssS*. Fragments containing strong intrinsic terminators (wild-type termination efficiency ≥75%^32^) are shown in pink. Below det., fragment below detection limit after selection (Methods). **e**, Enrichment of fragments tiling the genomic region antisense to *cssS*. Segment previously shown to induce Rho termination^6^ indicated by shaded purple region. Position indicates genomic coordinate. Fragments extending beyond plot boundaries are cut off by zigzag line.

Our massively parallel assay successfully distinguished genomic fragments that contain termination signals. While most fragments were depleted after chloramphenicol treatment, a distinct subpopulation (14%) was enriched by more than 10-fold. This subpopulation includes most annotated intrinsic terminators^32^ (Fig. 1b-d), whose enrichment values were correlated with previously estimated efficiencies, ranging from 65–99% termination (Extended Data Fig. 1).

Nearly half of the enriched fragments in the library were terminated by Rho, as evidenced by their sensitivity to bicyclomycin-benzoate (Fig. 1c, d; Methods). This subset of ∼10,000 fragments not only captured known targets of Rho termination but also resolved the sites of termination given the tiling nature of the library (Fig. 1d, e). Notably, Rho-terminated fragments are depleted of coding-strand sequences (1% fully overlap with genes vs. 29% in starting library; Fig. 2a), supporting our hypothesis that Rho termination signals are broadly excluded from mRNA sequence in *B. subtilis*. This cis-encoded selectivity contrasts sharply with what has been observed for *E. coli* Rho, which often terminates transcription of coding regions when translation is inhibited^1,5,8,9,11–13,22,24,26–30^.

**Fig. 2:**
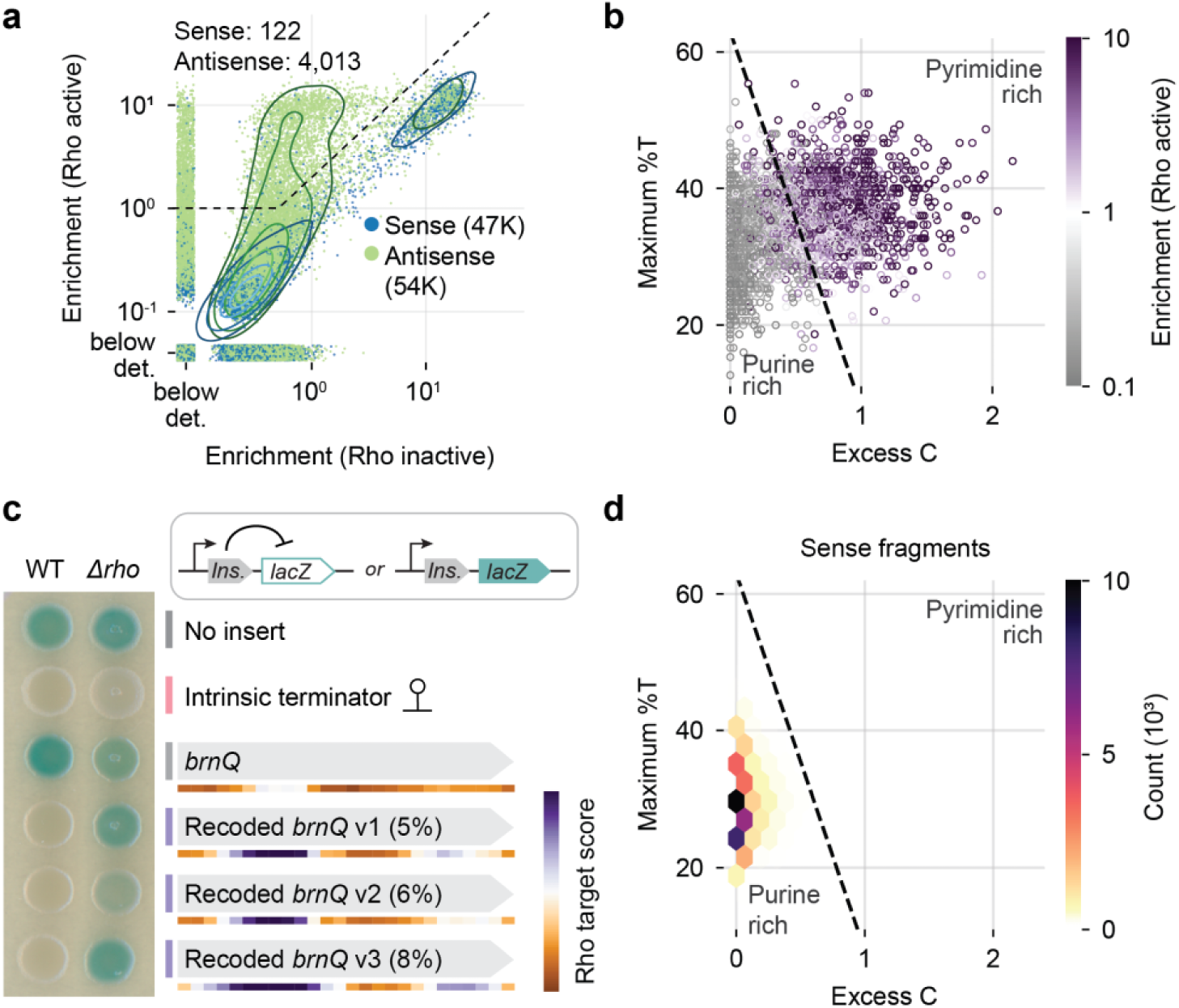
*B. subtilis* Rho terminates pyrimidine-rich antisense sequence. **a**, As in Fig. 1d, but for fragments that fully overlap with the sense (blue) or antisense (green) strand of annotated genes. Legend indicates total number of each fragment type. Top left, count of each fragment type in Rho-terminated fragments. Contour lines mark 10^th^, 25^th^, 50^th^, 75^th^ percentiles of Gaussian kernel density estimates, excluding points below detection. **b**, Sequence features of 1,500 nonterminated fragments (>500 reads pre-selection) and 1,500 Rho-terminated fragments colored by enrichment in the absence of BCM (fragment length ≥ 150 bp). Excess C quantifies C content relative to G content. Maximum %T is maximum T content in 150 nt windows tiling the sequence (Methods). Dashed line indicates the predicted boundary for Rho-terminated sequences (Rho target score = 10^0^) (Methods). **c**, Beta-galactosidase (LacZ) assays for termination of recoded *brnQ* variants in wild-type and Δ*rho* backgrounds. Top right, reporter schematic. Ins., insert; *lacZ,* beta-galactosidase gene. Left, image of *B. subtilis* reporter strains with indicated insert spotted on an X-gal plate (Methods). Percent of nucleotides substituted indicated in parentheses (insert sequences will be available in Supplementary Table 2). Heatmaps below each *brnQ* sequence show the Rho target score in 300-nt sliding windows (Methods). Colorbar range: 0.2 to 5 over log scale. Colored lines summarize result: grey, no termination; purple, Rho termination; pink, Rho-independent termination. Uncropped plate image in Extended Data Fig. 4a. **d**, Distribution of sequence features of the sense fragments shown in **a**, excluding fragments shorter than 150 nt (n = 46,777 fragments) (Methods). Dashed line as in **b**.

## Specificity of Rho-dependent termination

Only 5% of all genomic fragments (Fig. 1d) (7% of antisense fragments, Fig. 2a) are terminated by Rho, further illustrating the high degree of sequence specificity in *B. subtilis* Rho’s target selection. We identified two distinguishing features of Rho-terminated fragments. First, many of them possess long stretches (>150 nt) of high cytosine (C) content relative to guanine (G) (Extended Data Fig. 2a-c, Methods), consistent with previous observations of *E. coli* Rho’s sequence preference^8,14,20,23,25,33–35^. We developed a metric for excess C content that quantifies the cumulative CG bias in each fragment (Methods). Although fragments with high excess C values are nearly all terminated by Rho, those with intermediate values exhibit a wide range of termination activities (Fig. 2b and Extended Data Fig. 2d). We identified a second feature— thymine (T)-richness— that distinguished Rho-terminated fragments with intermediate excess C scores (Fig. 2b and Extended Data Fig. 2e). Together, a linear combination of these two pyrimidine-favoring metrics (“Rho target score”) separates the fragments by their termination activity, explaining 56% of the variability in enrichment (Fig. 2b and Extended Data Fig. 3).

To demonstrate that the presence of pyrimidine-rich sequence features is sufficient to induce Rho termination in *B. subtilis,* we increased the Rho target score of a native gene that is not regulated by Rho (*brnQ*) by introducing synonymous mutations. All three recoded versions (5-8% mutation rate) suppressed expression of a downstream *lacZ* included as a transcriptional reporter (Fig. 2c and Extended Data Fig. 4a). Deleting *rho* restored *lacZ* expression, indicating that recoding made these sequences targets of Rho-dependent termination (Fig. 2c). This result shows that the introduction of sequence signals recognized by Rho is sufficient to drive termination in *B. subtilis*.

Several lines of evidence indicate that the sequence requirement for *B. subtilis* Rho to terminate its RNAP is more stringent than in *E. coli.* First, a well-established target of Rho termination in *E. coli, lacZ*^22,27,28,36,37^, is not subject to Rho-dependent termination in *B. subtilis* (Fig. 2c, no insert control; Methods). Second, sequences that are known to be terminated by *E. coli* Rho in vitro^34^ do not have strong Rho target scores as we defined for *B. subtilis* (Extended Data Fig. 5a, b). Together, these results suggest that Rho termination occurs on a less diverse set of sequences in *B. subtilis* than in *E. coli*.

## Rho avoidance drives gene purine bias

Unlike the genes of other organisms, such as *E. coli*, we found that the coding strand of *B. subtilis* genes has strongly biased purine versus pyrimidine content (54% vs 46%, Fig. 2d and 3a). We hypothesized that this nucleotide bias is a result of selective pressure to avoid premature Rho termination of runaway RNAP, analogous to the depletion of termination factor motifs in eukaryotic genes^38,39^. By the rules of base pairing, the complementary pyrimidine bias is then present in antisense transcripts (Fig. 3a), priming their suppression by Rho. Our hypothesis predicts that in the absence of Rho, the gene purine bias would be lost during evolution. Conveniently, several isolated Bacilli lineages have lost *rho*^40,41^ (Fig. 3b), and these *rho*-less species indeed exhibit a much weaker purine bias in their coding sequences (Fig. 3c). Further, genes from *rho*-less species are prematurely terminated by Rho when expressed in *B. subtilis* (Fig. 3d and Extended Data Fig. 4b), confirming that the relaxed purine bias is accessible only in the absence of Rho. Therefore, for most Bacilli, the necessity to avoid Rho termination of runaway transcription likely drove the genome-wide separation between the nucleotide composition of sense and antisense strands.

**Fig. 3:**
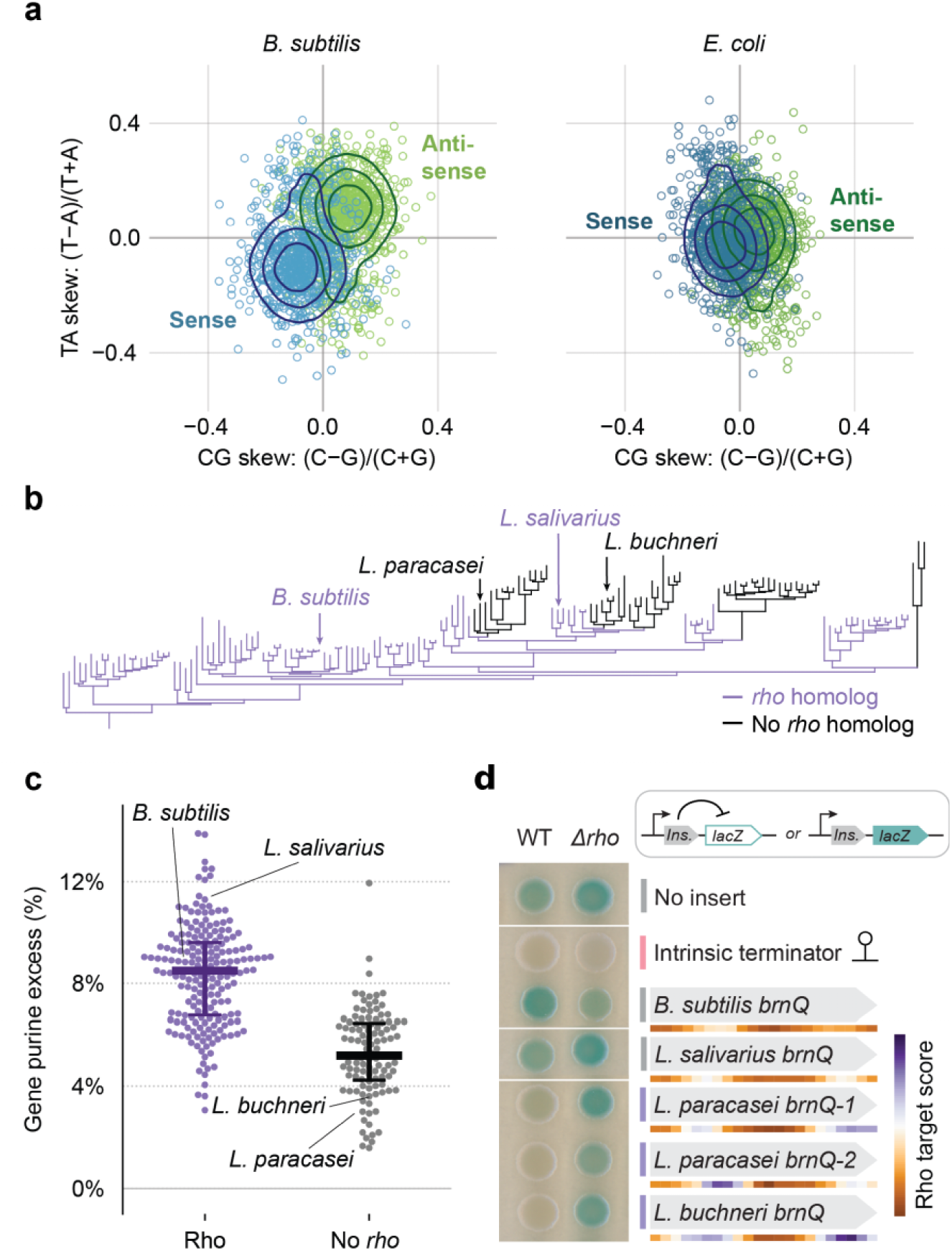
Rho avoidance drives purine bias in Bacilli genes. **a**, Nucleotide skew of 300-nt fragments of the *B. subtilis* (left) and *E. coli* (right) genomes derived from sense (blue) or antisense (green) strands of genes (Methods). Each point represents a single fragment. Contour lines mark 25^th^, 50^th^, and 75^th^ percentiles of Gaussian kernel density estimate. **b**, Phylogenetic tree of selected Bacilli species (155 species) colored by presence (purple) or absence (black) of a *rho* homolog in the genome^40^ (Methods). **c**, Average excess purine content in leading strand genes of 309 Bacilli species, separated by the presence or absence of *rho* in the genome (Methods). Purine excess quantifies the difference between purine and pyrimidine counts, normalized to length. Median, 25^th^, 75^th^ percentiles marked by horizontal lines. **d**, Beta-galactosidase (LacZ) assays for termination of *brnQ* homologs in wild-type and Δ*rho B. subtilis,* as in Fig. 2c. Colored bars summarize result: grey, no termination; purple, Rho termination; pink, Rho-independent termination. Insert sequences will be available in Supplementary Table 2. Uncropped plate image in Extended Data Fig. 4b.

We next addressed a competing hypothesis that the purine bias arose from mutational biases in DNA replication. Bacillota genomes have purine-rich leading strands of replication and pyrimidine-rich lagging strands, which has been attributed to their possession of dedicated DNA polymerases for synthesis in each direction (PolC and DnaE)^42–44^. As most genes reside on the leading strand (74% in *B. subtilis*^44^), replication-based nucleotide bias would result in an overall purine bias in coding sequences. To decouple replication biases from the Rho-based selective pressure we propose, we analyzed the purine-pyrimidine bias in lagging strand genes (Fig. 4a). Despite the overall depletion of purines on the lagging strand, coding sequences on this strand are purine-rich, with the median purine content closely matching that of leading-strand genes (53% vs. 54%, respectively) (Fig. 4b). This and other evidence^45^ suggest that purine bias is a feature of genes, not the direction of DNA replication, consistent with a need to avoid Rho-dependent transcription termination.

**Fig. 4:**
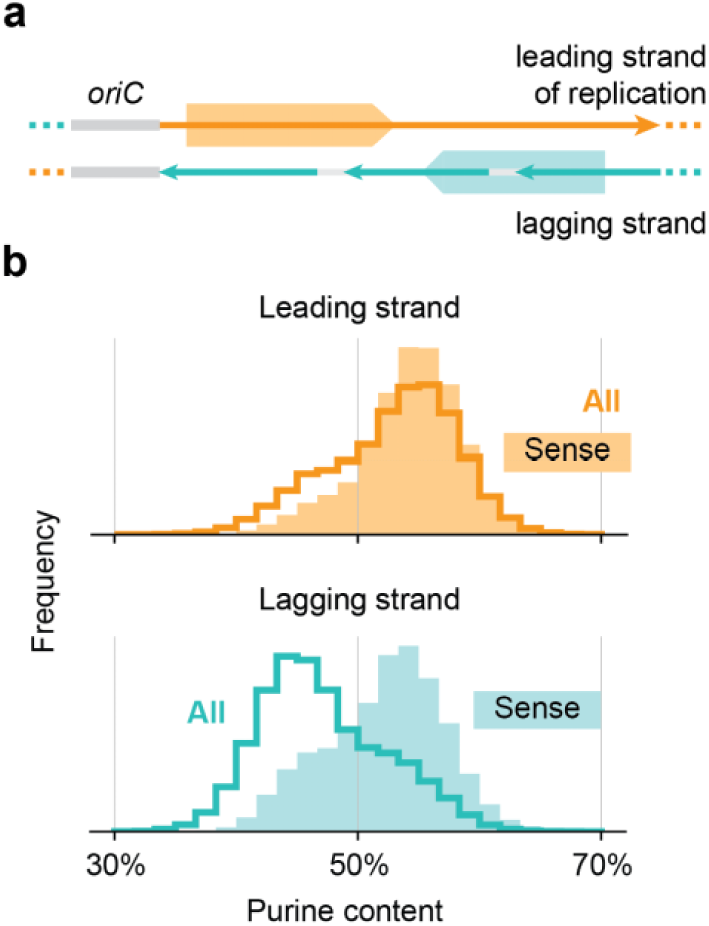
Purine skew is driven by the direction of transcription, not replication. **a**, Schematic illustrating regions considered in **b**. Thin arrows indicate direction of replication. Thick arrows represent coding sequences positioned on each replication strand. *oriC*, origin of replication. **b**, Top, purine content of 300-nt sequence fragments derived from any position on the leading strand (all; thick orange line), or sense to genes (sense; filled) (Methods). Bottom, same as top, but for the lagging strand of replication.

## Purine bias illuminates a hidden code

Purine bias has the potential to break the redundancy in the genetic code, as many amino acids can be encoded by synonymous codons of differing purine content. We compared the codon usage for these amino acids between Bacilli species that have maintained and lost *rho*. Focusing on the amino acids with four codons, where a purine-rich codon can be selected without necessitating a change in GC content, we found that codons that end with a purine are clearly favored in species that encode Rho (Fig. 5). Furthermore, some amino acids only have codons with an identical number of purines but can be replaced by other similar amino acids to increase the purine content^46^. For several such pairs of permissive substitutions, we observed a preference towards purine-rich amino acids in Bacilli species with Rho (e.g., glutamate with GAR codons vs. aspartate with GAY codons), although these preferences may reflect multiple constraints on amino acid usage (Extended Data Fig. 6). These results demonstrate that the purine bias creates a hidden layer constraining the usage of the genetic code.

**Fig. 5:**
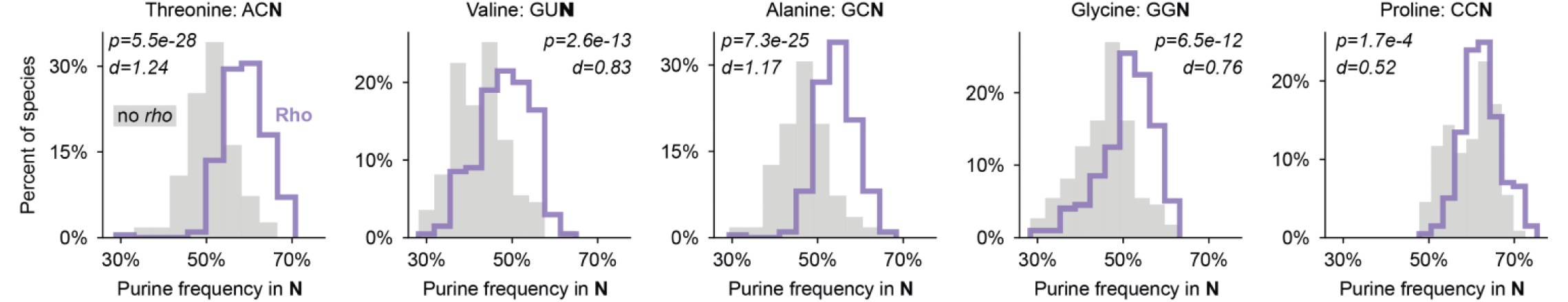
Rho shapes codon usage in Bacilli. Distribution of purine frequency in the wobble position among the codons used to encode the given amino acid across Bacilli species that do (thick purple lines; n = 200 species) and do not (filled grey bars; n = 111 species) encode *rho*^40^ (Methods). The five amino acids that are encoded by four codons are shown. N, any nucleotide. P-values from one-sided Mann-Whitney U tests. d indicates Cohen’s d effect size (Methods).

The hidden code imposed by Rho further creates a barrier for the expression of foreign genetic elements, such as bacteriophages and engineered constructs. To illustrate this barrier in the context of recombinant protein production, we show that transcription of a gene native to a different host (human growth hormone, HGH) is terminated by Rho in *B. subtilis* (Fig. 6a and Extended Data Fig. 4b). Expression can be restored upon synonymous recoding to increase purine content. This example demonstrates that foreign sequences that are not optimized to avoid Rho may not be compatible with runaway transcription in *B. subtilis.* This barrier could constrain horizontal gene transfer, suppress heterologous protein expression, and protect the host from selfish elements.

**Fig. 6:**
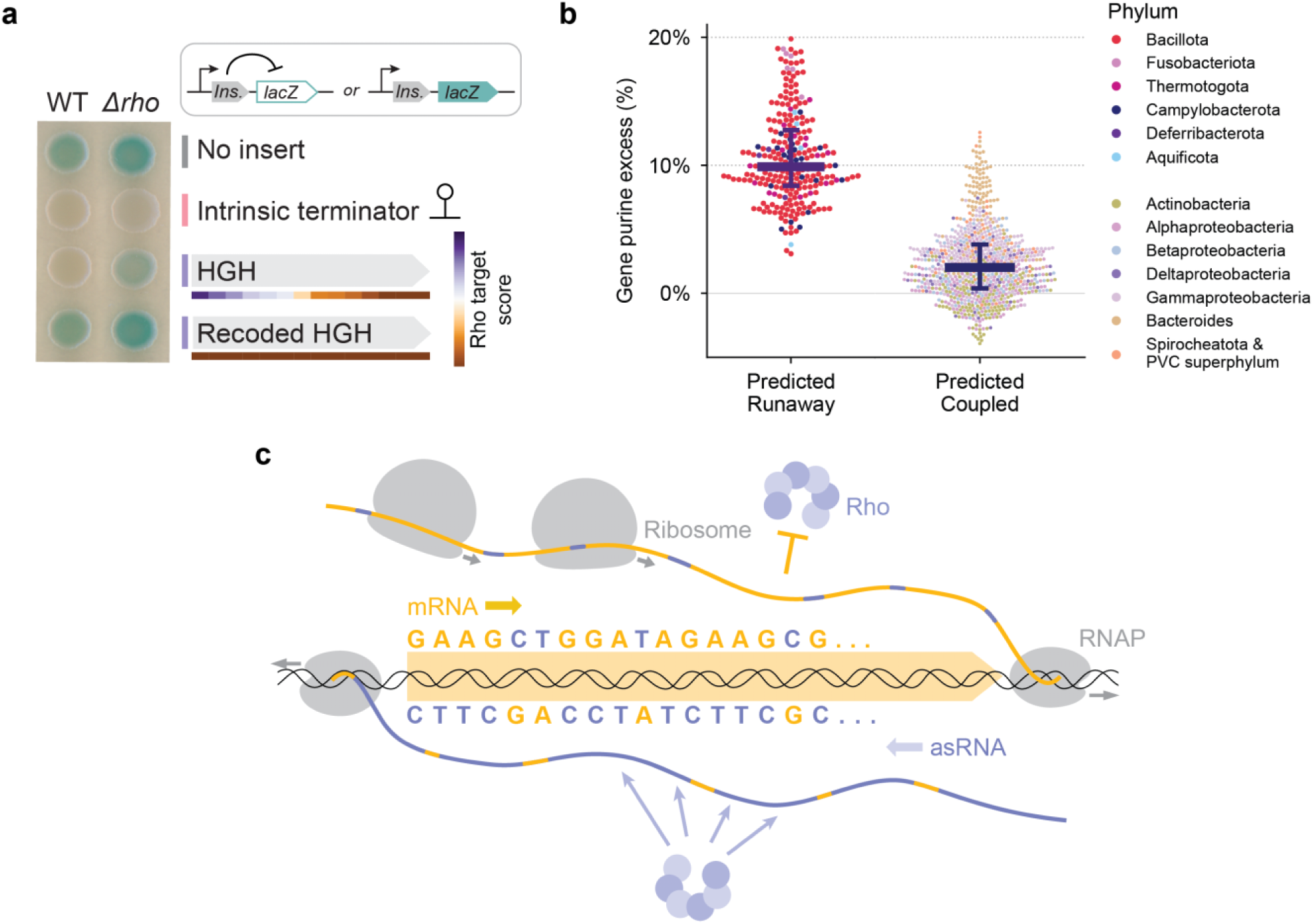
Purine-rich coding sequences protect runaway RNAP from Rho. **a**, Beta-galactosidase (LacZ) assays for termination of native and recoded human growth hormone (HGH) coding sequence in wild-type and Δ*rho B. subtilis,* as in Fig. 2c. Colored bars summarize result: grey, no termination; purple, Rho termination; pink, Rho-independent termination. Insert sequences will be available in Supplementary Table 2. Uncropped plate image is available in Extended Data Fig. 4b. **b**, Average excess purine content in leading strand gene sequences across n = 1,364 bacterial genomes that encode Rho, separated by phylum-level prediction of runaway or coupled transcription^32^ (Methods). Purine excess quantifies the difference between purine and pyrimidine counts, normalized to length (Methods). Median, 25^th^, 75^th^ percentiles marked by horizontal lines. **c**, Schematic of sequence-based selection of Rho targets during runaway transcription. Purine-rich genes are protected by Rho’s pyrimidine preference, while base pairing dictates that the antisense strand is enriched for pyrimidines, promoting Rho termination.

Beyond Bacilli, this hidden code likely applies broadly to species across the bacterial tree that have runaway transcription, including both gram-positive and gram-negative bacteria. In contrast to species predicted to have coupled transcription-translation, the coding genomes of species predicted to exhibit runaway transcription^6^ are consistently biased towards purines (Fig. 6b). The strong prevalence of purine-rich coding sequence in these species suggests that evolutionarily distinct phyla depend on a common sequence-based protection to avoid premature Rho termination, have similar constraints on mRNA and protein sequences, and represent an alternative paradigm for transcriptional surveillance.

## Discussion

We discovered that many bacterial genomes have evolved widespread purine bias in their coding sequences to escape termination during runaway transcription. This nucleotide code underpins the biased implementation of the genetic code, dramatically affecting gene expression levels and the fate of exogenous genetic elements. Meanwhile, antisense transcripts have an opposing pyrimidine bias that coincides with Rho’s heightened specificity, enabling Rho to maintain its function as a surveillance factor against spurious transcription^15,16^ (Fig. 6c). The broad separation of purines and pyrimidines into the coding and noncoding strands constitutes a new axis that shapes bacterial genomes, orthogonal to the well-appreciated GC-AT content.

The increased sequence specificity of Rho termination in *B. subtilis* compared to *E. coli* could be due to several factors, as this process involves both Rho and RNAP. First, the runaway RNAP in *B. subtilis* is nearly twice as fast as the *E. coli* RNAP^6,7^. A faster elongating RNAP could kinetically outcompete Rho, increasing the barrier to Rho termination^10–13,47^. Notably, *B. subtilis* RNAP pauses on T-rich sequences^48^, which could explain the increased Rho termination we observed for regions with high T content. Second, several trans-acting factors, including the transcription elongation factors NusG and NusA, nucleoid associated proteins, and direct Rho inhibitors, are known to modulate Rho activity in *E. coli*^10,12,14,20,21,37^. Deviations in the function of these proteins between *E. coli* and *B. subtilis* may also contribute to the differences in Rho specificity^48,49^. Finally, Rho termination requires both RNA-binding and ATP-dependent translocation^10–13,20,21^. In *B. subtilis,* either of these activities may have a stronger pyrimidine preference. Precisely mapping the exact positions of termination could help dissect how sequence influences each step of termination.

Additional processes may also influence the purine content of genes. Even in the absence of runaway transcription or Rho, genes in many organisms still exhibit a mild purine bias, including *E. coli* (Fig. 3a) and Bacilli that have lost Rho (Fig. 3c). In *E. coli*, transient uncoupling of ribosomes may make the RNAP prone to Rho termination^5,50^, which could be mitigated by reducing the pyrimidine content. More broadly, any mutational biases in DNA replication could also contribute to a weak nucleotide preference in gene sequences^51,52^. Indeed, although *B. subtilis* genes on both the leading and lagging strands of replication exhibit a purine bias to counteract Rho termination, the bias is stronger on the leading strand (Fig. 4b). Finally, requirements for amino acids with certain chemical properties may influence purine content through the sequences of the corresponding codons. When runaway transcription arose, a mild pre-existing purine bias from any of these factors might have set the stage for co-evolution of Rho’s heightened pyrimidine requirement and coding sequence composition.

Our study highlights that optimization of gene sequence is not merely defined by the effect of codon usage on translation—genes must adhere to requirements imposed by every step of gene expression. In bacteria, we show that a single transcription factor drives a profound coding constraint. We anticipate that systematic dissection of other central dogma processes will further unlock the principles for optimizing protein coding sequences in evolution and engineering.

## Extended Data Figures

**Extended Data Fig. 1:**
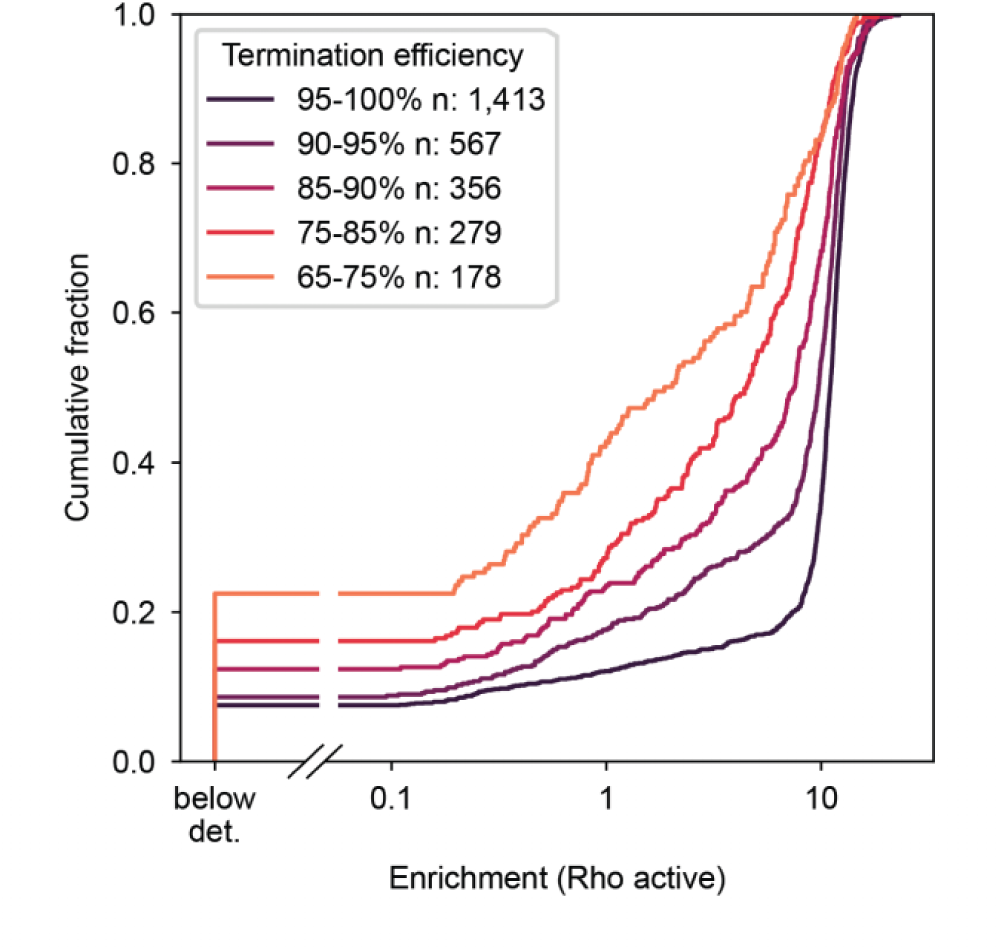
Fragment enrichment is correlated with termination efficiency. Related to Fig. 1. Cumulative distribution of enrichment in the absence of BCM (Rho active) for fragments containing an annotated intrinsic terminator, separated by termination efficiency^32^ (Methods). Fragments with more than 100 reads in the pre-selection sample are included. n indicates number of fragments in each distribution. Below det., below detection limit (Methods).

**Extended Data Fig. 2:**
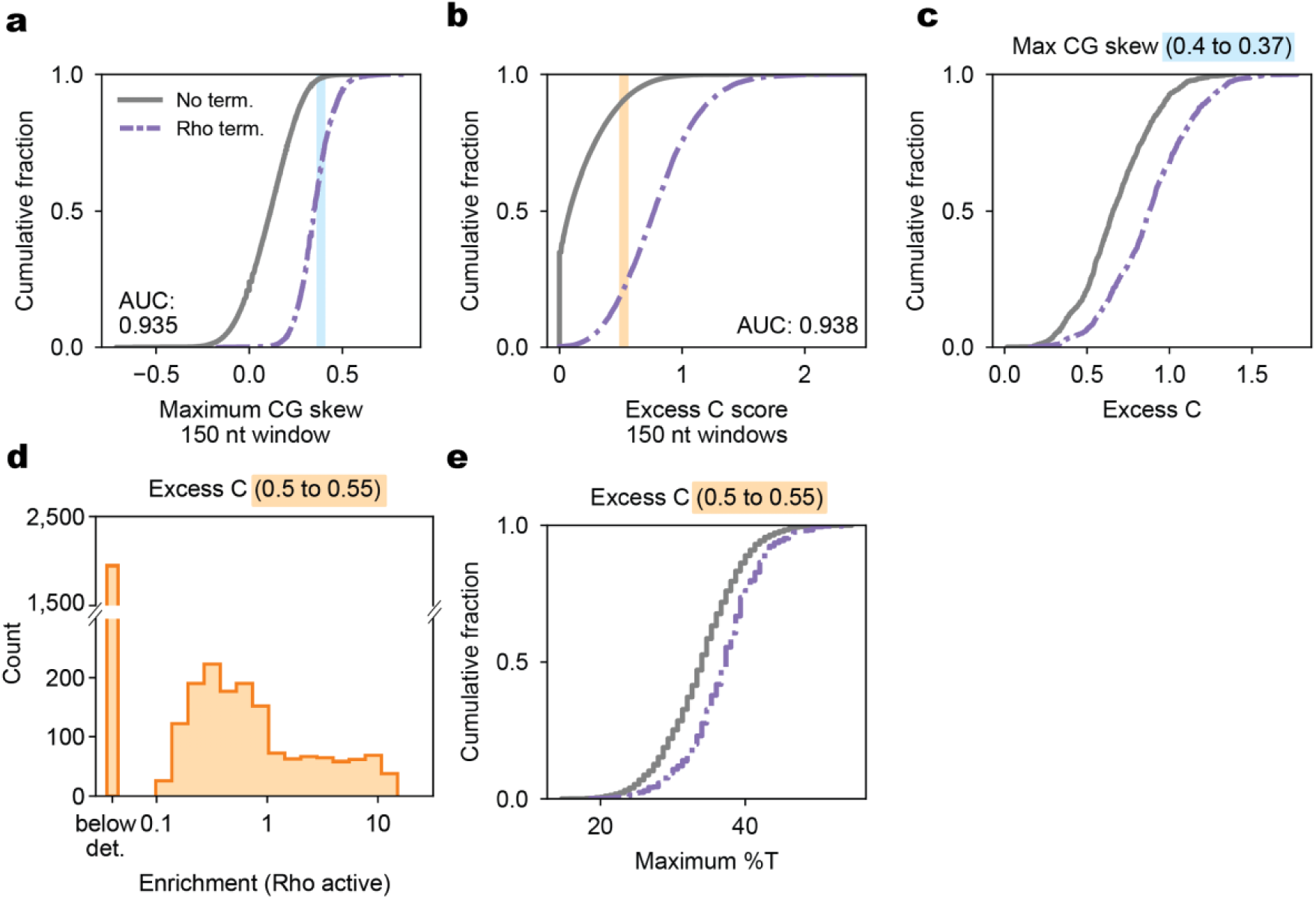
Performance of excess C and maximum %T scores. Related to Fig. 2. **a**, Cumulative distribution of maximum CG skew in 150 nt windows, separated by fragments (length ≥ 150 bp) with (dotted purple line; n = 9,983) and without (solid grey line; n = 136,728) Rho termination activity. AUC, area under ROC curve (Methods). Light blue shaded region indicates maximum CG skews considered in **c. b**, As in **a**, but for the excess C score, which is the sum of all positive CG skews in 150-nt windows tiling the fragment (Methods). Orange shaded region indicates excess C score of fragments shown in **d** and **e**. **c**, As in **b**, for fragments with a maximum CG score between 0.37 and 0.40 (region highlighted in blue in **a**). Non-terminated and Rho-terminated fragments with very similar maximum CG skews are separated by their excess C scores. **d**, Distribution of enrichments for fragments (length ≥ 150 bp) with an excess C score between 0.50 and 0.55. Below det., below detection (Methods). **e**, Cumulative distribution of maximum %T score for fragments with an excess C score between 0.50 and 0.55, separated by fragments with (dotted purple line; n = 9,983) and without (solid grey line; n = 136,728) Rho termination activity. Fragments with very similar excess C scores are separated by their maximum %T score.

**Extended Data Fig. 3:**
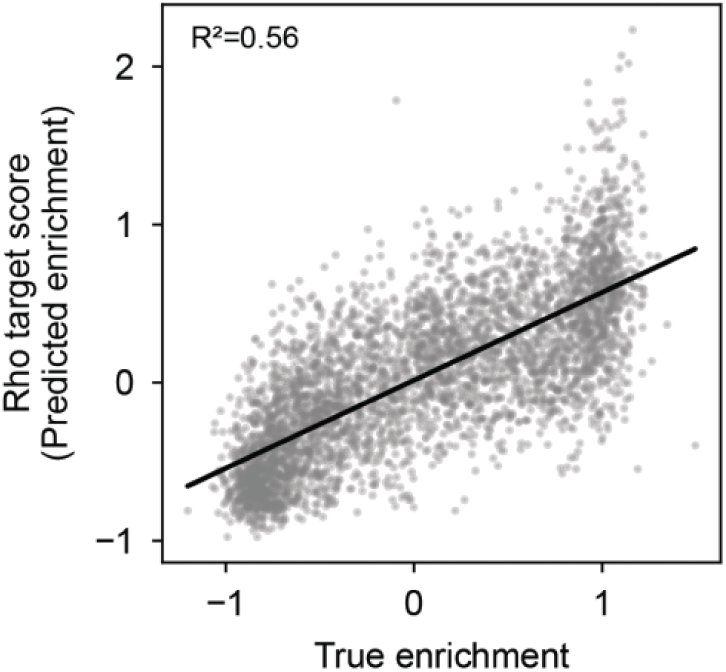
Multiple linear regression predicts enrichment from excess C and maximum %T scores. Related to Fig. 2, 3, 6. Scatterplot of predicted enrichment (Rho target score) vs. experimental enrichment for fragments retained in the test set (n = 3,958 fragments). Black line, linear regression between predicted and observed enrichment values (Methods). Enrichment was predicted by a linear model trained on the excess C and maximum %T values of 15,832 fragments split evenly into Rho-terminated and non-terminated fragments. All fragments in the test and training set were at least 150 nt in length. Pearson’s R^2^ value for the regression line shown at top left.

**Extended Data Fig. 4:**
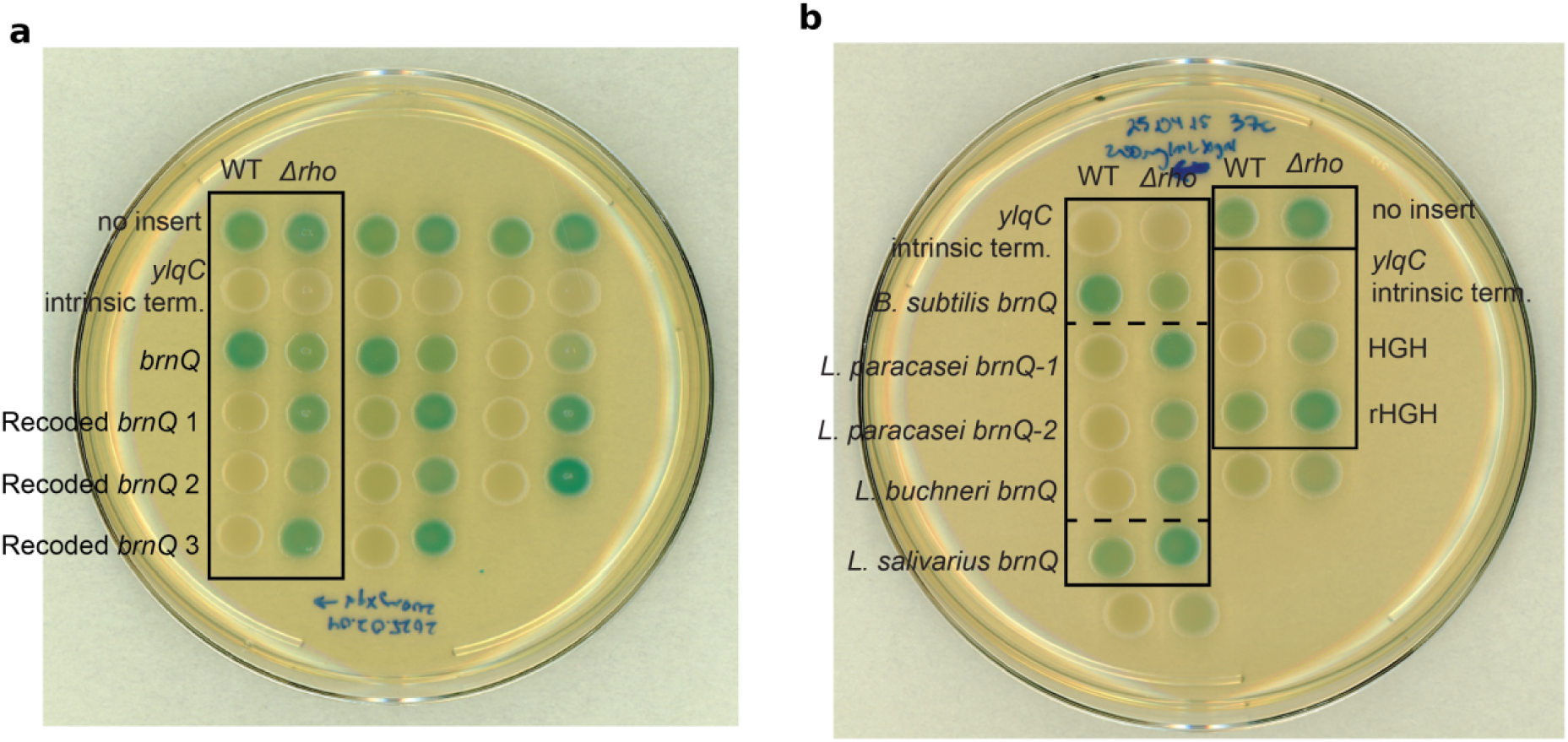
Uncropped X-gal plate images for LacZ assays. Related to Fig. 2, 3, 6. **a**, Full image for Fig. 2c. Black rectangle indicates cropping region. **b** Full image for Fig. 3d, 6a. Black rectangle indicates cropping region.

**Extended Data Fig. 5:**
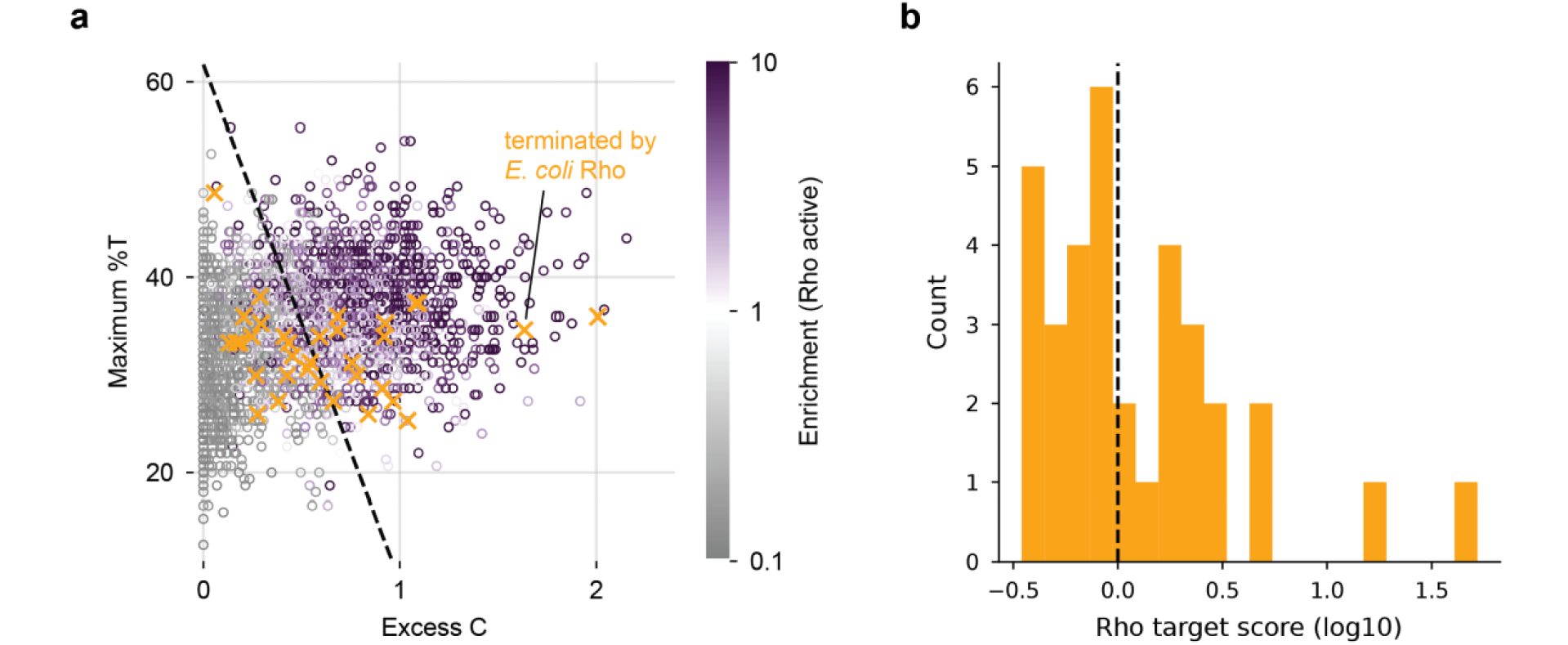
*B. subtilis* Rho is more selective than *E. coli* Rho. Related to Fig. 2. **a**, As in Fig. 2b, but overlaid with sequences strongly terminated by *E. coli* Rho *in vitro* (orange X)^34^. **b**, Distribution of Rho target scores for *E. coli* Rho-terminated sequences shown in **a**. Dashed line indicates enrichment cutoff of 1.

**Extended Data Fig. 6:**
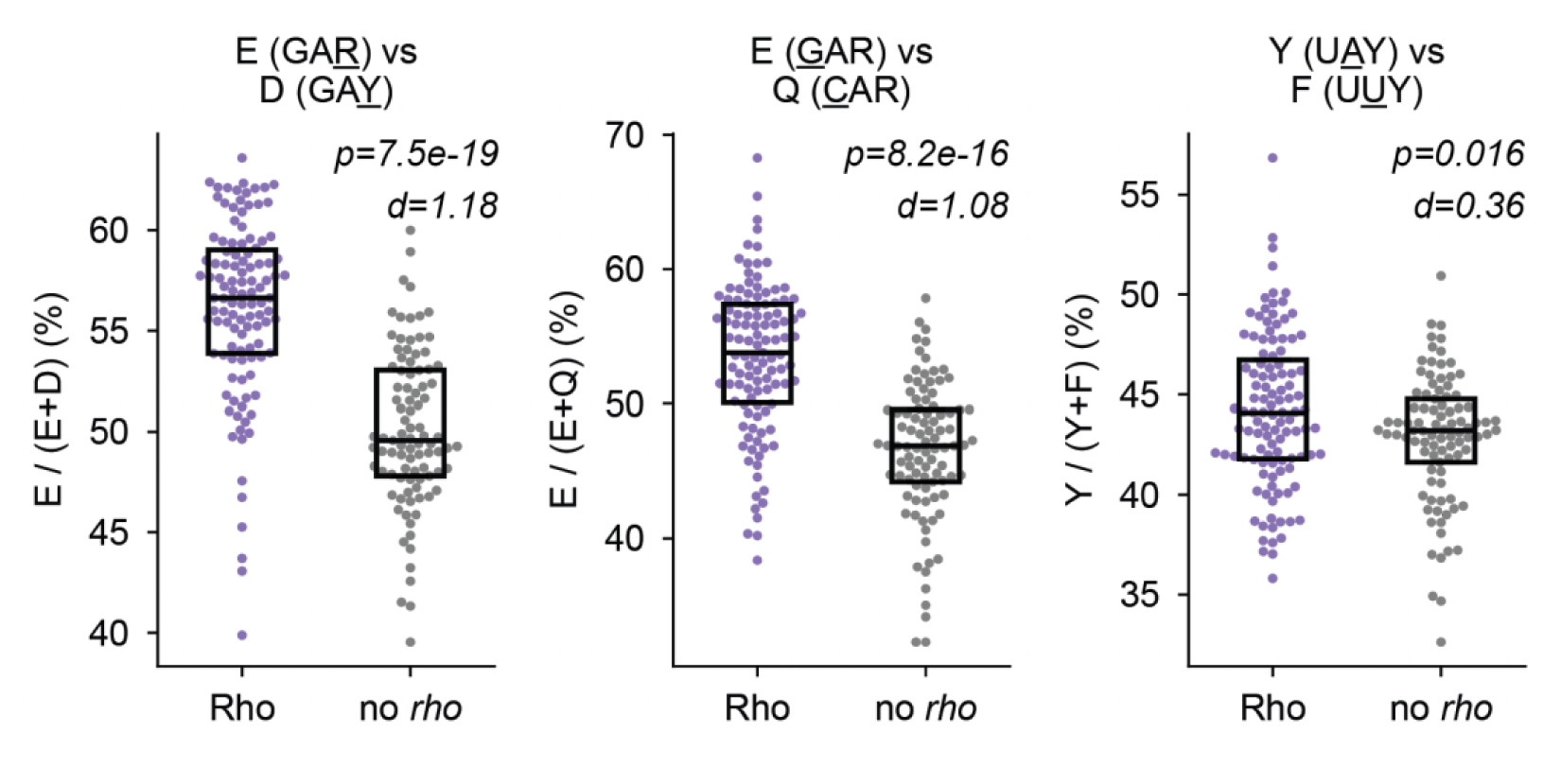
Rho shapes amino acid usage in Bacilli. Related to Fig. 3. Distribution of relative amino acid usage in Bacilli species separated by the presence or absence of a *rho* homolog^40^. Relative amino acid usage is shown for amino acid pairs with a strong (≥2) BLOSUM-62 substitution score^46^, where amino acid substitution requires a change in purine content in the nucleotide sequence without necessitating a change in GC content. Bacilli species shown have similar GC content (35–45%). n = 204 species (111 Rho, 93 no *rho*). Boxplots show median, 25^th^, 75^th^ percentiles. Underlined residue indicates position where purine content differs between amino acids. R, purine; Y, pyrimidine; N, any nucleotide. P-values from one-sided Mann-Whitney U tests. d indicates Cohen’s d effect size (Methods).

## Materials and Methods

### Strains

*B. subtilis* strains were generated from 168 (bearing the *trpC2* mutation) and *E. coli* gDNA was extracted from MG1655. Linearized plasmids and gDNA were transformed using either standard protocols relying on natural competence^53^or using supercompetent *B. subtilis* strains bJD086 (wild-type) and bJD087 (*Δrho*::*Kan*) based on^54^. bJD086 and bJD087 contain the *comK* induction system from SCK6^54^ inserted at *amyE*.

Sequences of plasmids and integrations were confirmed by Sanger or nanopore sequencing. All strains are listed in Supplementary Table 1. All plasmids were generated in *E. coli* DH5-α cells using standard protocols and are listed with additional details in Supplementary Table 1.

### Genomes

The RefSeq^55^ entries for *B. subtilis* (NC_000964.3) and *E. coli* (NC_000913.3) were used to map the sequencing reads and generate genome fragments in silico, and the corresponding annotations were used to classify fragments overlapping with sequences sense or antisense to genes.

### List of oligonucleotides

Supplementary Table 1 contains the list of oligonucleotides used for strain construction and sequencing library generation.

### Supercompetent *B. subtilis* transformation

To clone individual *B. subtilis* strains and generate the *B. subtilis* reporter library, linearized plasmid was introduced into *B. subtilis* using an inducible competence system mediated by the xylose-inducible *comK* allele from^54^ integrated at the *amyE* locus. Briefly, a colony of bJD086 (wild-type background) or bJD087 (Δ*rho* background) was picked into LB and grown to OD_600_ 0.8–1.1 at 37°C with vigorous shaking. At this point, 1% xylose (w/v) was added to induce expression of ComK. Induced cultures were shaken at 37°C for an additional 1.25–1.5 h, at which point 200 ng linearized plasmid was added per 100 μL induced cells. The cells were cultured with DNA for an additional 1–1.5 h before plating on selective plates for overnight growth at 37°C.

### Reporter plasmid library construction

The plasmid library was constructed through isothermal assembly of tagmented gDNA fragments into the reporter plasmid backbone, pJD16. To generate the inserts, a 1:1 mixture of *B. subtilis* and *E. coli* gDNA was fragmented by Nextera XT tagmentation (Illumina). 6 ng of the tagmented gDNA was PCR amplified with oJD173/oJD174 for 10 cycles using Q5 DNA polymerase (New England Biolabs). PCR products 200–400 bp in length were size-selected using an 8% TBE polyacrylamide gel (ThermoFischer). This size selected DNA was then cleaned up with a DNA Clean & Concentrator-5 column (Zymo Research). To linearize the pJD16 backbone, pJD16 plasmid was PCR amplified with oJD091/oJD092 using Q5 DNA polymerase. The PCR product was extracted from an agarose gel using a Zymoclean Gel DNA Recovery Kit (Zymo Research) and DpnI (New England Biolabs) digested for 60 minutes at 37°C. The DpnI-treated DNA was cleaned up with a DNA Clean & Concentrator-5 column.

To construct the plasmid pool, 200 ng of the pJD16 backbone PCR was mixed with insert at a 1:4 molar ratio and assembled at 50°C for 15 min in a Gibson Assembly reaction (New England Biolabs). This reaction was then cleaned up with a DNA Clean & Concentrator-5 column and transformed into electrocompetent *E. coli* (New England Biolabs 10-β cells). Transformants were plated on LB with 100 μg/mL carbenicillin in 245 mm square bioassay dishes (Corning) and grown at 37°C overnight. Cells were then scraped off the plates for ZymoPure II Maxiprep plasmid extraction (Zymo research).

### Transformation and integration of reporter library into *B. subtilis* genome

The supercompetent transformation protocol described above was used to integrate the reporter library into *B. subtilis* strain bJD086, replacing the *comK* induction cassette at the *amyE* locus. Briefly, a colony of bJD086 was picked into 22 mL LB and induced with 1% xylose (w/v) once the culture reached OD_600_ 1.02. After 1.5 h of growth in xylose, 15 mL of induced culture was combined with 1.5 mL (30 μg) of plasmid pool linearized with ScaI-HF (New England Biolabs). This digest was prepared per the manufacturer’s protocol with 1 μg plasmid per 50 μL reaction. After 1.5 h of incubation with DNA, cells were pelleted, resuspended in residual media, and plated on 245 mm bioassay dishes containing 100 μg/mL spectinomycin. After growth overnight, cells were scraped into LB with 100 μg/mL spectinomycin and approximately 500 million cells were back-diluted into LB with 100 μg/mL spectinomycin. This culture was grown for 5 h and 1 mL aliquots of culture were mixed 1:1 with 40% glycerol and frozen at −80°C.

### Library growth and harvesting

To perform chloramphenicol selection on the *B. subtilis* library to enrich for genomic fragments with transcription termination activity, the library was grown in LB for 5 generations, split into +BCM and DMSO control (−BCM) conditions for pre-treatment, and then back-diluted into selective media containing chloramphenicol and BCM or chloramphenicol and DMSO.

To begin the initial outgrowth, a single glycerol stock was thawed, resuspended in warm LB, and back-diluted to OD_600_ 0.006 in 50 mL warm LB. This starter culture was grown at 37°C with vigorous shaking to OD_600_ 0.2. A second back-dilution to OD_600_ 0.006 in 25 mL was then performed to pre-treat the cultures with bicyclomycin-benzoate (BCM) or a DMSO control. For this pre-treatment, cells were grown either in LB with 120 ng/mL BCM (+BCM treatment) or 2.5% DMSO (−BCM treatment). The pre-treatment cultures were then grown to OD_600_ 0.2, at which point both cultures were harvested (pre-selection harvest). At the end of the +BCM pre-treatment, a small amount (3%) of residual chloramphenicol acetyltransferase is expected to remain in cells with inserts driving Rho termination, which may explain the slight increase in +BCM enrichment for these fragments relative to the non-terminated fragments (Fig. 1d). BCM concentration and pre-treatment duration were optimized using the *kinB* Rho termination site as a positive control.

To begin the chloramphenicol selection, the BCM pre-treated culture (+BCM) was back-diluted to OD_600_ 0.013 in 25 mL of LB with 5 μg/mL chloramphenicol and 120 ng/mL BCM, and the untreated culture (−BCM) was back-diluted to OD_600_ 0.0075 in 25 mL of LB with 5 μg/mL chloramphenicol and 2.5% DMSO. The cultures were then grown with vigorous shaking at 37°C for 2.75 h before the post-selection harvest (OD_600_ 0.24–0.27). To harvest cells, 11–14 mL of culture was pelleted at 4°C and then frozen in liquid nitrogen. Cell pellets were stored at −80°C until gDNA extraction. In total, cell pellets for four conditions were collected: +BCM culture before and after chloramphenicol selection; −BCM culture before and after chloramphenicol selection.

### gDNA sequencing

Sequencing libraries were prepared to quantify the frequency of each genomic fragment variant in the four experimental conditions. gDNA was extracted from the cell pellets using the Wizard Genomic DNA Purification kit (Promega) following manufacturer’s instructions. To reduce RNA contamination, the gDNA was then cleaned up with the genomic DNA clean and concentrator-10 (Zymo Research). Libraries were generated using a two-step Q5 PCR protocol. To attach UMIs, a 2 cycle PCR was performed using 2 μg of gDNA per 150 μL PCR reaction (scaled as needed for library complexity) with primers oJD180/oJD182. This PCR reaction was cleaned up with two rounds of magnetic bead clean-up at a 1:1 ratio of beads to sample (PCRClean DX bead, Aline Biosciences). Half (pre-selection libraries) or one quarter (post-selection libraries) of this cleaned-up reaction was then amplified in a second PCR with primers oJD181/oJD183. The second PCR reaction was gel-purified using an 8% TBE gel. These libraries were sequenced with 50 bp paired-end reads on a G4 sequencer (Singular Genomics).

### Fragment quantification

Quantification of genomic fragments in each sample was handled using custom Python scripts. First, the paired-end sequencing reads were aligned to the *B. subtilis* (NC_000964.3) genome using the bowtie package (version 1.2.3)^56^. 40 nt of each read was used for alignment, and alignments with more than 1 mismatch or an insert size of more than 5,000 bp were discarded (bowtie arguments: -trim3 10 -v 1 -X 5000). Any two reads with the same aligned start and end position and UMI were collapsed to a single read. The number of unique reads mapping to each genomic fragment was calculated and normalized to the total number of reads in the sample. The enrichment of each fragment in each condition (+BCM or −BCM) is the ratio of its frequency in the post-selection sample to the corresponding pre-selection sample. Raw read counts for all fragments and samples will be available in Supplementary Table 3.

### Thresholding and pseudocounting enrichments

To reduce noise in the enrichment values, a threshold of ≥50 reads per fragment was applied to the pre-selection samples and a threshold of ≥25 reads per fragment was applied to the post-selection samples. Fragments that fell below the post-selection read threshold due to depletion during selection were assigned a pseudo-enrichment value of 0.015 (labeled as below detection in Fig. 1d, 2a, and Extended Data Fig. 1, 2d) provided that their raw enrichment was below 0.5 (−BCM) or 1 (+BCM). 161,256 fragments passed these thresholds or were assigned a pseudo-enrichment in both conditions (+BCM and −BCM), and their reported enrichments will be available in the Fig. 1 Source Data. In Fig. 1d and 2a, fragments with a pseudo-counted enrichment were assigned a random enrichment value to jitter points for visualization. These fragments are annotated in the Source Data.

### Assignment of Rho-terminated and non-terminated fragments

Fragments terminated by Rho should be enriched in the absence of BCM, when Rho is active, and this enrichment should be lost when Rho is inactivated by BCM treatment. Two criteria were used to identify these fragments from their enrichment scores. First, Rho-terminated fragments had to exceed an enrichment of 1 in the −BCM condition, which roughly corresponds to ≥65% termination efficiency (Extended Data Fig. 1). Second, the enrichment in the +BCM condition had to be more than 2-fold lower (0.45x) than the enrichment in −BCM. This cutoff was chosen to minimize misclassification of fragments with known intrinsic terminators as Rho-dependent termination sites (less than 1.6% misclassified). 9,995 fragments were classified as driving Rho-dependent termination activity, the positions of which will be available in the Source Data for Fig. 1 and Supplementary Table 3. Non-terminated fragments were defined as those with an enrichment less than 1 in the −BCM condition.

### Fragment overlap with sense and antisense regions and intrinisic terminators

Fragments overlapping with sense and antisense regions of the *B. subtilis* genome were identified using the annotated RefSeq CDS features for NC_000964.3. Sense and antisense fragments were required to fully overlap with the same or opposite strand of an annotated CDS, and any fragments that overlapped with multiple coding regions (on either strand) were discarded. 47,550 sense and 54,692 antisense fragments of *B. subtilis* are shown in Fig. 2a. Fragments were annotated as containing an intrinsic terminator if any of the wild-type positions of intrinsic termination identified in^32^ fell within the fragment (n = 3,037 fragments). The termination efficiencies used in Extended Data Fig. 1 were determined from the readthrough fraction in Rend-seq of Δ*pnpA B. subtilis*^32^.

### Excess C and maximum %T metrics

To identify sequences with a skewed cytosine (C) content relative to guanine (G), we wanted to capture information about the magnitude of the difference between C and G counts in a defined region (representing a single Rho termination site) and also capture the cumulative effect of multiple regions where C content exceeds G content, which we hypothesized would increase the probability of Rho termination.

When we considered magnitude alone by identifying the highest CG skew ((C−G)/(C+G)) in windows tiling the sequence fragments, we found that a relatively short window (150 nt) best distinguished the Rho-terminated fragments (Extended Data Fig. 2a); averaging over longer sequences appeared to dilute the signal. We found that combining CG skew across multiple windows further improved the prediction of Rho-terminated fragments. To capture the contribution of multiple windows with high CG skew, we summed the CG skew across all 150-nt tiling windows (step size: 50 nt) with positive CG skew (the number of Cs exceeds Gs). We refer to this metric as the “excess C” score (Extended Data Fig. 2b). A score of 0 indicates that in all 150-nt windows, the number of Gs matched or exceeded the number of Cs. We found that the cumulative information in the excess C score was important for predicting Rho termination: Rho-terminated and non-terminated sequences with similar maximum CG skews could be further differentiated by their excess C scores (Extended Data Fig. 2c). The window length, tiling step size, and skew cutoff were optimized for the highest predictive power in differentiating Rho-terminated and non-terminated fragments.

Maximum %T was determined by counting the number of threonines (Ts) in every 150-nt window tiling a sequence fragment and then taking the maximum T count as a fraction of the window length. We found that the performance of metrics related to T content in distinguishing the Rho-terminated fragments was not improved by incorporating information about relative adenine content. Fragments shorter than 150 bp (2.1% of input library) were excluded from sequence feature analysis.

### Linear model and Rho target score

To evaluate the explanatory power of the excess C and maximum %T metrics and predict enrichments in silico, a linear model was trained to predict fragment enrichments (log_10_-transformed) from these two scores. Fragments with enrichments below the detection limit were excluded from the test and training sets. The model was trained on 15,832 fragments (length ≥ 150 nt) split evenly into Rho-terminated and non-terminated fragments and the variance explained (Pearson R^2^ value) was generated by evaluating the model on a reserved test set of 3,958 fragments (Extended Data Fig. 3). Fragments overlapping with 5′ ends identified by Rend-seq^32^ were excluded to avoid confounding effects from promoter activity. The coefficients of the model are 0.0208 for maximum %T and 1.144 for the excess C score, and the intercept is −1.31. This linear model was also used to predict enrichments in silico from sequences’ excess C and maximum %T scores, and these predicted enrichments are defined here as the “Rho target score.” The R value reported in the text and Extended Data Fig. 3 corresponds to the Pearson’s correlation coefficient between the observed and predicted enrichments in the test set (n = 3,958 fragments).

The boundary line shown in Fig. 2a, c illustrates the positions on the plot where this Rho target score is 1 (log_10_ -transformed enrichment of 0). Using our linear model, sequences that fall to the left of this line would be predicted to be depleted in our experiment, and sequences to the right would be predicted to be enriched due to Rho termination. The equation of the line is: maximum %T = −55.0 (excess C score) + 63.0.

### LacZ reporter strains

DNA fragments with potential Rho-dependent transcription termination sites were cloned upstream of *lacZ,* under the control of the IPTG-inducible pSpankHy promoter in the vector pJD19. To generate pJD19, a restriction enzyme cloning site was introduced to pGJ02^6^ and *lacZ-α* was replaced by a full-length *lacZ* allele optimized for expression in *B. subtilis. lacZ* was amplified from pESW830 (from E. Wirachman, *lacZ* sequence originally derived from pCAL1422^57^). This version of *lacZ* aligns to residues 45–3,075 of the *E. coli lacZ* sequence with 4 mismatches (full sequence is available in^6^). As *lacZ* Rho termination sites were identified between residues 140 and 425 in *E. coli*^36^, we do not expect that the missing N-terminal sequence could explain the lack of termination in *B. subtilis*.

Recoded *brnQ* variants, native and recoded HGH sequences, and *brnQ* homologs were synthesized as Gene fragments (Twist Biosciences) and amplified by PCR, adding the native *brnQ* ribosome binding site (15 bp upstream of start codon) to drive translation. Wild-type *B. subtilis brnQ* and the *ylqC* intrinsic terminator control were amplified by PCR from gDNA. These PCR products were inserted into pJD19 by isothermal assembly or digested alongside pJD19 with EagI-HF and HindIII-HF (New England Biolabs) before ligation using QuickLigase (New England Biolabs). The resulting plasmids were then linearized by ScaI-HF (New England Biolabs) digestion and transformed into the *amyE* locus to replace the *comK* expression system in bJD086 (wild-type background) and bJD087 (*Δrho::Kan* background) as previously described. For strains expressing wild-type or recoded versions of *B. subtilis brnQ,* the native copy of *brnQ* gene was then knocked out through transformation of gDNA from the *ΔbrnQ::erm* strain following natural competence protocols.

For qualitative assays of β-galactosidase activity, blue-white screening was performed on X-gal (5-Bromo-4-chloro-3-indolyl β-D-galactopyranoside) (GoldBio) plates. For each strain, a colony was picked into 5 mL LB and grown at 37°C with vigorous shaking to OD 0.50–3.5. These cultures were then diluted to OD_600_ 0.005 in LB and 3 μL of the dilutions were spotted onto LB-agar plates with 1 mM IPTG and 200 μg/mL X-gal. Plates were incubated at 37°C overnight and imaged 16–17 h after plating.

### Recoding *brnQ* and HGH

To increase the Rho target score of *brnQ,* synonymous mutations were introduced as follows. First, the codons corresponding to each amino acid were ranked first by the number of cytosine (C) and uracil (U) bases, and then by the number of adenine (A) bases. Codons with the highest C+U and A content were ranked highest so that codons with high pyrimidine content and low G content would be prioritized in the recoding. The three rare codons in *B. subtilis* (CUA, AUA, AGG)^58^ were not included in the ranking. A subset of the residues in *brnQ* were then randomly selected for recoding. For each of these residues, the wild-type codon was replaced by the codon with the highest ranking for the corresponding amino acid (maintaining the wild-type codon if a higher rank could not be achieved). Recoded sequences with regions of high excess C and maximum %T content were chosen manually through comparison to the data in Fig. 2b. The sequence of each recoded variant will be available in Supplementary Table 2.

For recoding of human growth hormone (HGH), the native sequence encoding the mature HGH peptide (generated after cleavage of signal peptide) was acquired from RefSeq (NM_000515.5). A similar procedure was then followed to remove pyrimidine-rich features through synonymous mutation. For HGH, codons were ranked first by the number of cytosine (C) and uracil (U) bases, and then by the number of guanine (G) bases. Codons with the lowest C+U and highest G content were ranked highest, and rare codons were again removed. Recoding was performed as described above. A recoded sequence with minimal excess C and low maximum %T content was chosen manually through comparison to the data in Fig. 2b. The sequence of the native and recoded variants will be available in Supplementary Table 2.

### *brnQ* homologs in Rho and non-Rho species

*brnQ* homologs were identified by gene name in the RefSeq annotations for *Lacticaseibacillus paracasei* (NC_022112.1)*, Lentilactobacillus buchneri* (NZ_CP073066.1), and *Ligilactobacillus salivarius* (NZ_CP117983.1). Protein sequence homology was verified by EMBOSS needle alignment^59^: reported similarities ranged from 49–55%.

### Rho homolog classification

For *Lentilactobacillus buchneri* and the genomes shown in Fig. 6b, classifications of genomes with and without a *rho* homolog were taken from^32^. For all other Bacilli species (Fig. 3c-e, Extended Data Fig. 6), the Rho homolog classifications in^40^ were used.

### In silico generation of *B. subtilis* and *E. coli* genome fragments

To analyze the nucleotide composition of sense and antisense sequences in the *B. subtilis* and *E. coli* genomes, 300-nt tiling fragments (step size: 50 nt) were generated from the sequence of all CDS features in the RefSeq annotation files. Features shorter than 300 nt were excluded. The antisense fragments were generated by taking the reverse complement of the sense sequence fragments. All fragments (n = 51,531 for *B. subtilis* and 57,477 for *E. coli*) were used to generate the Gaussian kernel density estimates shown in Fig. 3a, and a subset of 1,600 fragments, split evenly into sense and antisense strand, are shown in the scatterplot. For the unbiased leading and lagging strand fragments shown in Fig. 4, randomly selected start positions were used to generate 300-nt windows of genomic sequence.

For Fig. 4, the *B. subtilis* sense sequence fragments and randomly generated fragments were assigned to the leading or lagging strand based on their position relative to the origin (4,215,389^60^), assuming replication arms of equal length.

### Bacilli phylogenetic tree

The phylogenetic tree of Bacilli (Fig. 3b) was generated from the GTDB tree^61^ (version 226, downloaded June 2025). 155 Bacilli species analyzed in^40^ and *L. buchneri* were included in the tree, with Rho homolog classification based on species name using^32,40^ as described above. Only a subset of the 309 Bacilli species shown in Fig. 3c were included in this tree for visualization purposes. See the Fig. 3 Source Data for the full list of species included in the tree.

### Sense strand purine content in Bacilli and across Bacteria

To compare the purine content in sense sequences across Bacteria (Fig. 6b), we analyzed a set of 1,551 RefSeq genomes with coupling predictions^32^ and an annotated origin of replication^62^. To correct for length variation across genes, the purine content was determined for a random 250-nt fragment of each annotated gene (genes shorter than 250 nt were discarded). Each gene was then assigned to the leading or lagging strand using the position of the origin of replication from^62^ and assuming replication arms of equal size. The average purine content was calculated separately for genes on the leading and lagging strand, and the leading strand averages for the 1,364 genomes that encode a *rho* homolog are shown in Fig. 6b. The designation of runaway vs. coupled transcription was performed at the phylum level, based on^32^. Species in Bacillota, Fusobacteria, Thermotogota, Deferribacterota, Aquificota, and Campylobacterota were classified as exhibiting runaway transcription. This data will be available in the Fig. 6 Source Data. The same process for analysis of sense sequence purine content was repeated for the 308 Bacilli genomes (GTDB classification of c Bacilli) from^40^ and *L. buchneri* to generate the data for Fig. 3c.

### Codon usage

For each of the Bacilli genomes (GTDB classification) with a *rho* homolog assignment from^40^, the nucleotide sequences of the annotated gene sequences (CDS features) were downloaded from the NCBI RefSeq database^55^. The frequency of each codon across the entire coding genome was then normalized to the amino acid frequency to determine the codon usage by amino acid.

### Overlap with *E. coli s*ense sequence

To identify fragments fully overlapping with sense sequence in *E. coli,* the process described in *Fragment overlap with sense and antisense regions and intrinisic terminators* was followed for the *E. coli* CDS annotations (NC_000913.3).

### Relative amino acid usage

For each of the Bacilli genomes (GTDB classification) with a *rho* homolog assignment from^40^, the nucleotide sequences of the annotated CDSs were downloaded from the NCBI RefSeq database^55^. To control for differences in GC content, a cohort of 204 species with similar GC content (35-45%) is shown. The frequency of each amino acid in the entire coding genome was determined as the sum of its codon frequencies. For amino acid pairs with a BLOSUM-62 score of 2 or greater^46^, where substitution requires a change in purine content without necessitating a change in GC content (E to D, E to Q, Y to F), the frequency of the purine-rich amino acid was divided by the total frequency of both amino acids in the pair.

### Statistics

To generate the p-values reported in Fig. 5 and Extended Fig. 6, a one-sided Mann-Whitney U test was performed. Assuming underlying distributions of the same shape, the alternative hypothesis for the test was that the median relative frequency (of codon or amino acid usage, respectively) was higher for Bacilli with *rho* than species without *rho.* The values for the U statistic are as follows: Fig. 5: T (19,385), V (16,585), A (18,872), G (16,243), P (13,820); Extended Fig. 6: E to D (8,853), E to Q (8,507), Y to F (6,056). Cohen’s d was used to report the effect size of the difference in means (absolute difference between the means divided by pooled standard deviation) and is reported in the corresponding figure. The number of each species is available in the figure legend, and source data will be provided.

For Extended Data Fig. 2a-b, the AUC statistic was computed for the distribution of the maximum CG skew or the excess C score, respectively, in non-terminated fragments (n = 136,728 fragments) vs. Rho-terminated fragments (n = 9,983 fragments). Fragments shorter than 150 nt were excluded from the analysis. Source data will be provided.

A description of the calculation of the R^2^ statistic reported in the text and Extended Data Fig. 3 is available above, in the section *Linear model and Rho target score*.

## Data availability

All data generated or analysed during this study are included in this published article (and its supplementary information files). Source data for each figure and extended data figure will be included in the published article.

## Code availability

The Python (version 3.11.7) scripts used to calculate fragment enrichments (calculate_enrichment_pc.py), assign termination status (assign_Rho_termination.py), calculate sequence feature metrics (excessC_maxT_scores.py), compute the Rho target score (calculate_rho_target_score.py), and calculate average purine excess in bacterial coding genomes (gene_purine_excess.py), will be deposited on Github (jdierksheide/Rho) and be available on Code Ocean.

Other data analysis and plotting was performed using custom Python scripts that are available upon reasonable request from the authors.

## Acknowledgements

We thank A. Grossman’s laboratory for providing plasmids; BM Koo for providing *B. subtilis* single-gene deletion strains; A. Grossman and members of the G.-W.L. laboratory for discussions; J.-B. Lalanne, A. Savinov, J. Xue, J. Cascino, I. Kim, E. Ermis, and D. Bartel for comments on the manuscript; members of the BioMicroCenter at MIT for help in performing DNA sequencing. G.W.L. is an investigator at the Howard Hughes Medical Institute. This research was supported by NIH R35GM124732 (to G.W.L), the Smith Odyssey Award (to G.W.L.), MathWorks graduate fellowship (to K.J.D.), and NSF Graduate Research Fellowships (to J.C.T. and G.E.J.).

## Author contribution

Conceptualization of the massively parallel reporter assay for Rho-termination: GEJ and JCT; Conceptualization of other experiments and analyses: KJD, GWL; Methodology: KJD, JCT, GEJ; Investigation: KJD, JCT, GEJ; Visualization: KJD; Writing: KJD, GWL; Editing: JCT, GEJ; Resources: GWL; Supervision: GWL.

## Materials & correspondence

Correspondence and requests for materials should be addressed to Gene-Wei Li.

